# On the distribution and function of synaptic clusters in dendrites

**DOI:** 10.1101/029330

**Authors:** Romain D. Cazé, Amanda J. Foust, Claudia Clopath, Simon R. Schultz

## Abstract

Local non-linearities in dendrites render neuronal output dependent on the spatial distribution of synapses. A neuron will activate differently depending on whether active synapses are spatially clustered or dispersed. While this sensitivity can in principle expand neuronal computational capacity, it has thus far been employed in very few learning paradigms. To make use of this sensitivity, groups of correlated neurons need to make contact with distinct dendrites, and this requires a mechanism to ensure the correct distribution of synapses contacting from distinct ensembles. To address this problem, we introduce the requirement that on a short time scale, a pre-synaptic neuron makes a constant number of synapses with the same strength on a post-synaptic neuron. We find that this property enables clusters to distribute correctly and guarantees their functionality. Furthermore, we demonstrate that a change in the input statistics can reshape the spatial distribution of synapses. Finally, we show under which conditions clusters do not distribute correctly, e.g. when cross-talk between dendrites is too strong. As well as providing insight into potential biological mechanisms of learning, this work paves the way for new learning algorithms for artificial neural networks that exploit the spatial distribution of synapses.

## 1 Introduction

Active synapses distribute non-randomly on dendrites: synaptic contacts from nearby neurons have been observed to form clusters (Druckmann et al., 2014; Rah et al., 2015). Functional synaptic clustering has been observed. Takahashi et al. (2012) demonstrated that nearby synapses tend to activate more often during spontaneous activity in adult rats, and (Kleindienst et al., 2011) also reported synaptic clustering in organotypic slices from neonatal rats during development. Finally, Makino and Malinow (2011) reported that synaptic clustering can evolve during sensory experience. These reports leave open the functional role of such a non-random synaptic distribution.

Theoretical studies have predicted the existence of synaptic clusters (Mel, 1992; Poirazi and Mel, 2001; Poirazi et al., 2003). Because dendrites, even if they are passive (Koch et al., 1982), integrate non-linearly, the relative position of synapses exerts a salient influence on the neuronal input-output function. Therefore, a clustered synaptic spatial distribution can enhance the computational capacity of single neurons. In particular, clustered synaptic configurations allow neurons to compute linearly inseparable functions (Zador et al., 1993; Caze et al., 2012), can enhance auditory coincidence detection (Agmon-Snir et al., 1998) and can help to compute binocular disparity (Archie and Mel, 2000), as well as contribute to whisker directional tuning in the barrel cortex (Lavzin et al., 2012). All these interesting computational properties, however, arise when synaptic clusters distribute evenly on a neuron. Correlated ensembles of neurons need to generate distinct clusters, and a single ensemble should not predominate.

We present here a learning algorithm that guarantees a useful distribution of synaptic clusters. Several previously proposed learning algorithms are capable of generating synaptic clusters (Iannella and Tanaka, 2006; Wu and Mel, 2009; Legenstein and Maass, 2011). To guarantee that the same cluster does not form twice they gate learning using somatic or dendritic spiking. This strategy has at least one unavoidable constraint: the action potential needs to invade the entire dendritic arbor. It is known, however, that the somatic action potential can fail to reach terminal dendrites (Spruston et al., 1995; Vetter et al., 2012), which makes this an unsatisfactory solution. Here we instead guarantee the uniqueness of synaptic clusters by supposing that, on a short time scale, a presynaptic neuron makes a constant number of synapses on a postsynaptic neuron.

## Materials and Methods

### Controlling correlation in ensembles of input spikes

We used correlated Poisson processes to model the input spikes (Brette, 2009). Each process is the sum of an independent process with a frequency *f*(1−*c*) and of a process common to the ensemble with a frequency *f* × *c* (*f* in hertz and *c* ∈ [0,1] without units). Fig. 1A shows input spikes that were generated using this method. In this study, input spikes are grouped in seven ensembles, each composed of 100 neurons, with a mean firing rate of 10 Hz and 80% of their spikes correlated within the ensemble. These input spikes lasted 40s, made up of 4000 bins of 10ms. During the last 20s of each set of input spikes, we rotated the set of ensembles by 50.

**Figure 1:**
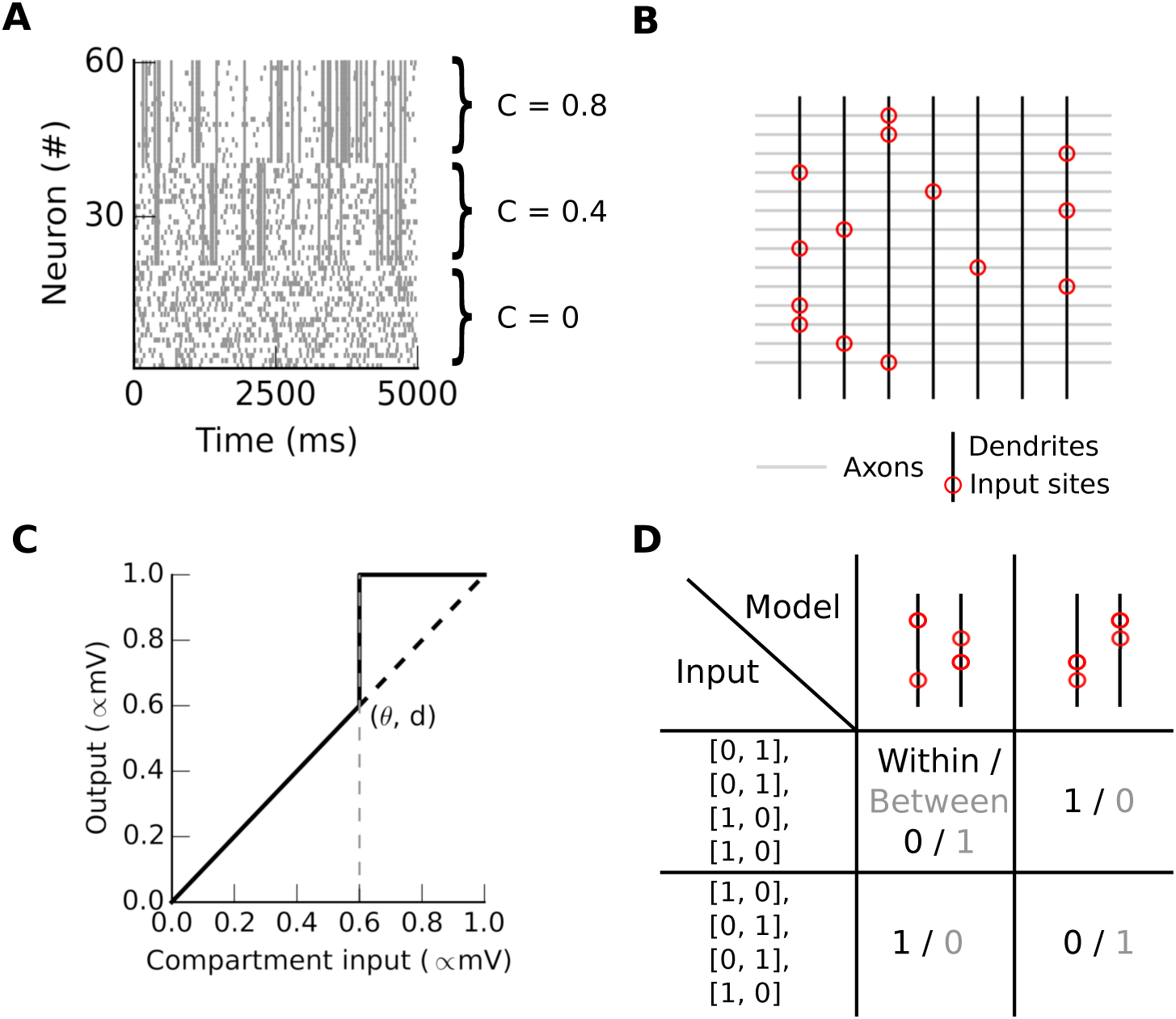
A multi-subunit model for studying the spatial distribution of synaptic clusters. **A**. Raster plot showing the activity of three ensembles, each with a different level of internal correlation. Neurons 0-19, 20-41 and 42-61 have correlation coefficients of 0, 0.4 and 0.8 respectively. **B** The input sites (red circles) are where the connections are most numerous on dendritic compartments (black vertical lines). **C** The non-linear transfer function of the dendritic compartment. The input is the weighted sum of the presynaptic activity, and the output of this function is transmitted to the soma. **D** Co-activation probability of a synapse pair either within (black) or between (gray) compartments given a synaptic architecture (column) vs a set of input spikes (row). There are four synapses coming from four neurons during two time steps.

### A multi-subunit binary neuron model

We computed the mean somatic depolarization in time bins of 10ms using our multisubunit model. This computation has two steps, as in (Poirazi et al., 2003). For each bin, the binary inputs *x_i,j_* first sum linearly in each compartment, given a local set of weights the *w_i,j_* compartment *j*. We present an example of synaptic architecture (set of synaptic weights) in Fig 1B. Secondly, we non-linearly transform the normalized sum resulting in *v_j_* = *D_j_*(∑_*i*_ (*w_i,j_x_i,j_*). *v_j_* denotes a local signal that could be interpreted as either the mean membrane potential or the mean calcium concentration in compartment *i*. *D* is a function with a threshold *θ* at which *v* jumps to 1. Then all the *v* are linearly summed at the soma to determine whether the neuron fires or not: *y* = *S*(∑_*j*_*v_j_*), where *S* is a Heaviside function with threshold Θ = 0.2. This value for Θ means that 20% of the compartments need to reach their maximum depolarization to elicit a somatic spike. We used here a somatic threshold of Θ = 0.2, a dendritic threshold of *θ* = 0.2, and synaptic weights bounded between 0.01 and 1 modeling only excitatory synapses.

In contrast with more detailed biophysical models, our model lacks temporal integration, and its compartments process inputs independently. We have, however, previously demonstrated that this model would nonetheless yield the same conclusions if we were to relax these assumptions (Cazé et al., 2013). This simplicity enables rapid simulations and in-depth analyses that would be harder in a detailed biophysical model.

### A local learning rule and horizontal normalization

The learning rule uses the local signal *v_j_* of the *j^th^* compartment to compute the weight change Δ*w_i,j_* = *α*(2*x_i_*−1)*v_j_*, where *α* = 0.1 is the learning rate. Consistent with experimental data, the learning rule depends only on a local signal in the dendrites. Plasticity depending only on dendritic spikes, independent of somatic spikes, has been observed experimentally on a number of occasions (Feldman, 2012; Mehta, 2004; Kim et al., 2015).

In each time bin, the synaptic weights arising from a given source are normalized by the total synaptic weight from this source. This is to account for the limited number of synapses per afferent (Branco and Staras, 2009). We introduced this normalization to foster an optimal distribution of synaptic clusters on dendrites (compare movie S1 and S3 respectively with and without normalization), and to focus on the spatial distribution of synapses rather than the total synaptic strength from an afferent.

### Co-activation probability

We define the co-activation probability as the number of times a pair of synapses are active, divided by the total number of times either synapse in the pair activates, analogous to previous experimental studies (Kleindienst et al., 2011; Takahashi et al., 2012). The co-activation probability depends on two distinct variables: the input spike and the set of synaptic weight values (see Fig. 1D). Here we illustrate a case with two dendrites receiving input at four synapses over two time steps. For instance, in the middle column and row, the inputs in both steps activate one synapse on each dendrite, giving a “between” co-activation probability of one. The “within” co-activation probability here is zero.

To compute the co-activation probability, we first select the compartment bearing the highest weight for a given input, indicated by empty circles in subsequent figures. We then either replay the input spikes, or replay the seven input vectors corresponding to the seven ensembles. In the latter case, each input vector corresponds to the activation of one ensemble. We then select every possible pair of synapses, and classify each as occurring either “within” or “between” compartments. The co-activation probability is the number of times a pair of synapses are active, divided by the total number of times either synapse in the pair activates.

## Results

### Correlated inputs distribute synapses in multiple distinct clusters

We show here that correlated inputs evoke synaptic clusters that distribute evenly on dendrites. First we examine a neuron receiving inputs from two ensembles and featuring two compartments as in Fig. 2. We describe the evolution of the two mean synaptic weights, one per ensemble, using a 2D vector field. Importantly, this 2D vector field demonstrates only two stable point: (0,1) and (1,0) in all conditions. Hence synapses either from the first or the second ensemble grow together to form a cluster (Fig. 2 upper left, “linear”). Saturation within a compartment when it activates more than 50% ((*θ* = 0.5, *d* = 0.5)) sharpens the vector field toward the stable points (Fig. 2 upper right, “non-linear”). Interestingly, horizontal normalization, in which each input has a constant total synaptic weight, guarantees that a cluster forms only once. Without horizontal normalization, the same ensemble can form clusters on multiple compartments.

**Figure 2:**
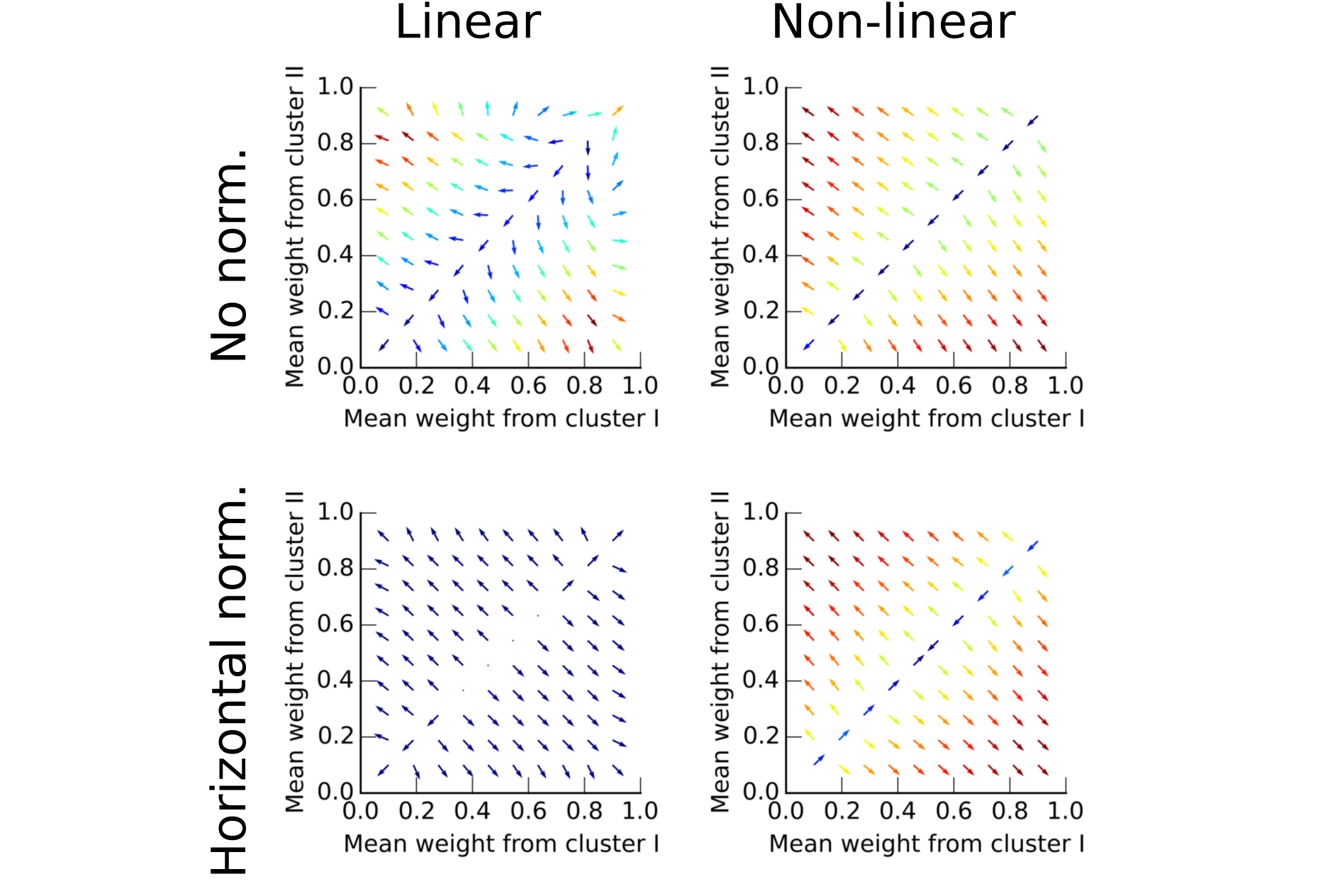
Vector fields showing the evolution of the mean synaptic weight per cluster. The arrow’s origin is the mean synaptic weight value. It points in the direction of the average evolution of these means and is color-coded given the intensity of this movement (blue:low, red:high). We used semi-analytic results to construct this figure. Note that only (0,1) and (1,0) stay stable in the four situations, with or without normalization and with linear (*θ* = 1, *d* = 1) or non-linear summation in compartments (*θ* = 0.5, *d* = 0.5).

We scaled up our implementation to show that seven ensembles, made of 100 neurons each, can form evenly distributed synaptic clusters (Fig. 3). Before receiving any input, synapses distribute randomly on the neuron’s seven subunits (Fig. 3A). Note that each dendrite (vertical black line) corresponds to only one subunit. The appearance of spatial clustering within each dendrite is due to the adjacent positioning of correlated inputs for visual clarity, but inputs could impinge anywhere on the compartment. After receiving inputs organized in seven correlated ensembles (Fig. 3C), seven synaptic clusters form on the neuron with one unique cluster per ensemble (Fig. 3B). Movie S1 illustrates that horizontal normalization gives rise to even cluster distribution. Without normalization, synaptic clusters form, but the same cluster can occur multiple times as illustrated by Movie S3.

**Figure 3:**
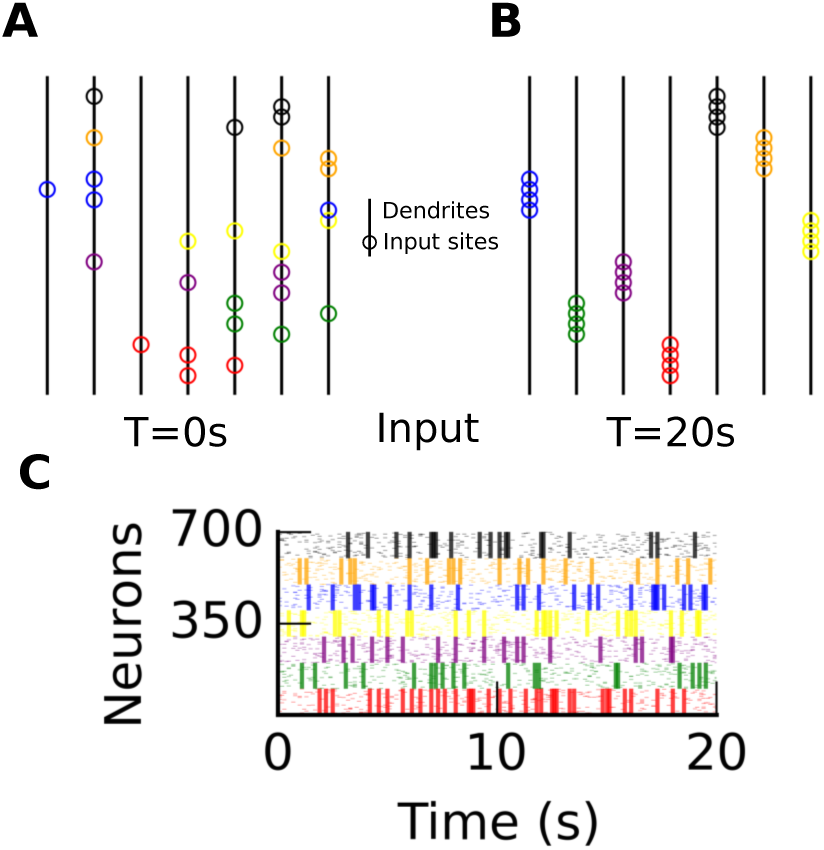
Correlated inputs generate synaptic clusters on dendrites. **A** and **B** A multi-subunit model where each subunit (vertical lines with *θ* = *d* = 0.14) before (T = 0s) and after learning (T = 20s). Circles indicate the most probable synaptic sites. Each color corresponds to a different correlated ensemble. **C** Input spikes (*n* = 700) grouped in seven correlated ensembles. Each ensemble has a distinct color, and 80% of spikes within an ensemble are correlated.

Non-linear integration in dendrites can render a neuron sensitive to scattered synaptic activation. For low dendritic threshold, a clustered learned input ensemble will trigger a single dendritic spike, whereas a scattered ensemble will trigger many. For a high somatic threshold, a neuron requires multiple dendritic spikes to drive a somatic spike. Because our learning rule gives rise to evenly distributed clusters on a model with non-linear compartments, a novel unlearned stimulus can saliently activate the neuron (Fig 4A).

**Figure 4:**
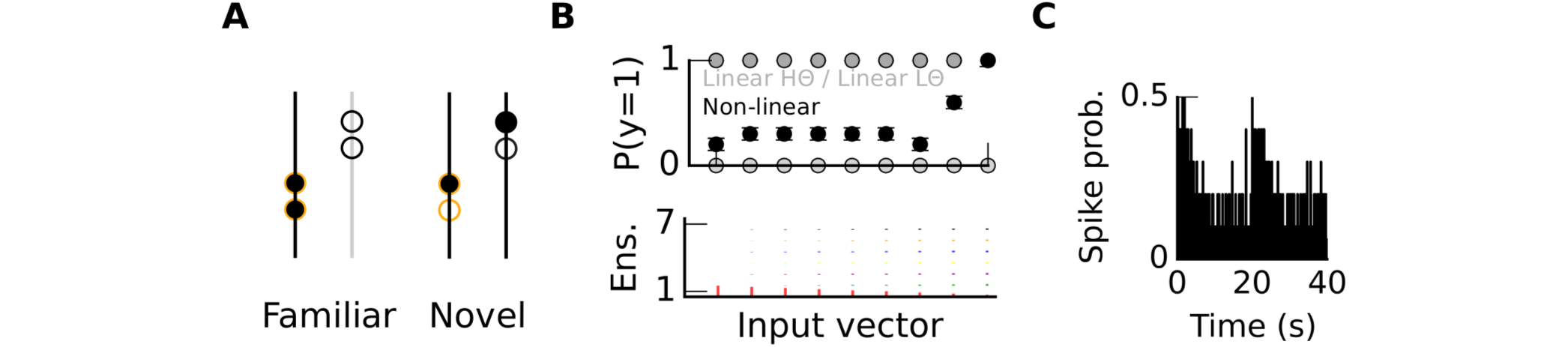
Familiar and novel inputs create distinct spatial distribution of synaptic activity on neurons. **A** Activity within two subunits (vertical lines) with four synaptic sites (circles) after learning. Activity distributes differently if the stimulus is familiar (experienced during learning) or novel (never experienced); activity is color-coded (white:inactive / black:active). **B** Mean spiking probability (over a hundred model instances) after learning for 9 different input vectors (each vector as a length of 700). The models feature either linear (gray) or non-linear (black *d* = *θ* = 0.12) subunits. The linear models can either have a low (light gray) or high (dark gray) somatic threshold. That is, fewer or more than a 100 active synapses are required to trigger an action potential, respectively. In the non-linear case, more than half of the subunits need to be active to trigger an action potential. **C** Mean spiking probability for each time bin during learning (average of over a hundred model instances). In the linear case this would have been a flat line because of horizontal normalization (see Methods).

The sensitivity to the spatial distribution of inputs could render the neuron sensitive to unexpected inputs. We measured the response of two types of neuron models integrating their inputs either linearly or non-linearly(100 instance of each model).

In the non-linear case the firing probability increases with the novelty of an input vector (Fig 4B black line). If the input vector corresponds to the activation of all inputs from a learned ensemble, then the post-synaptic neuron remains silent. If the vector corresponds to the activation of inputs from different ensembles, then the neuron fires. In summary, a neuron integrating its input non-linearly can signal an unfamiliar input deviating from the learned set of ensembles. This is illustrated by the mean firing probability plotted as a function of time (Fig 4C) where we present a new ensemble set at *t* = 20s.

In the linear case, however, the neuron responds to all input vectors equally because each source of input spikes makes a contact with a total synaptic weight always equal to one (see Methods). Therefore, for a somatic threshold larger than 100/700 (the size of an ensemble being *n* = 100), a linear neuron fires for all input vectors, and for a threshold lower than 100/700, a linear neuron stays silent for all input vectors (Fig 4B grey lines).

Next we examine how changes in input statistics affects the reshaping of synaptic clusters.

### Changes in input statistics can reshape synaptic clusters

We started from a model displaying synaptic clusters (*t* = 20*s* in Fig 5A), and then exposed it to inputs organized into a new set of ensembles between *t* = 20*s* and *t* = 40*s* (Fig 5C), shifting the set of ensembles by 50 neurons to obtain the new set of ensembles. For instance, while neurons 1 to 100 were correlated between 0s and 20s, after the rotation, neurons 51 to 151 are correlated. This rotation reshaped the spatial distribution of synapses to reflect the new set of ensembles (Fig 5B). Thus, long-lasting changes to input statistics reshape the synaptic architecture. In contrast to classical unsupervised algorithms, this learning occurs continuously in our model. These changes can be visualized in Movie S3 and S4, respectively, with or without horizontal normalization.

**Figure 5:**
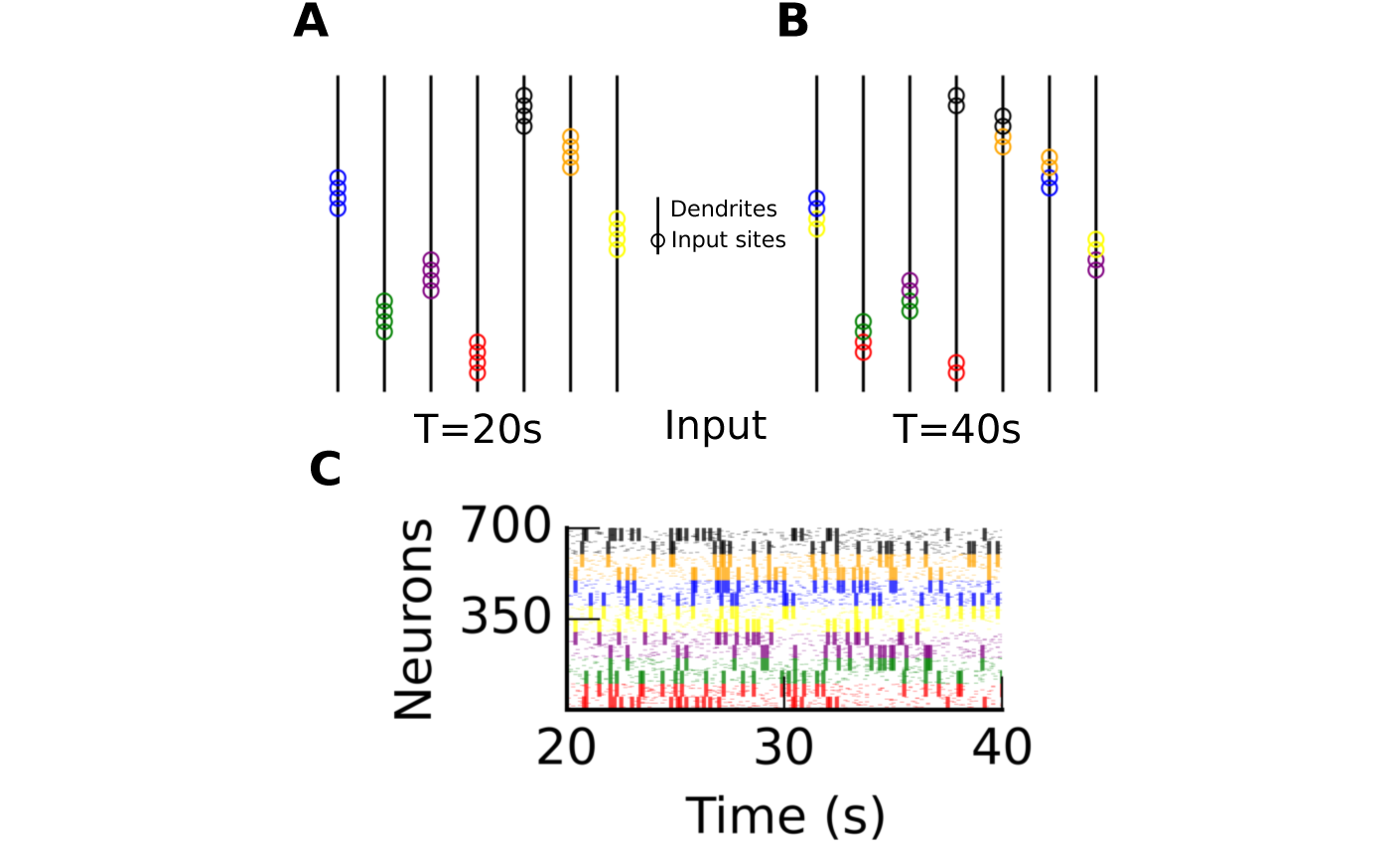
Changes in input statistics can reshape synaptic clusters. **A** and **B** A multi-subunit model at time *T* = 20*s* and *T* = 40*s*. All subunits (vertical lines) are non-linear (*θ* = *d* = 0.14). Empty circles indicate synaptic sites (color-coded given the ensemble set at *t* < 20*s*). **C** Input spikes between *t* = 20*s* and *t* = 40*s*.

### Co-activation probability quantifies the distribution and formation of synaptic clusters

We show that the formation of synaptic clusters in our model recapitulates co-activation probabilities measured in experiments (Kleindienst et al., 2011; Takahashi et al., 2012) (Fig. 6). The co-activation probability is the fraction of time bins in which both synapses activate (see Methods for examples). Kleindienst et al. reported that co-activation probability depends on the distance between spines (Fig. 6A). Similarly in our model, coactivation probability is higher for “within” compared to “between” subunits. ΔCo-ac is the difference between these two cases. Moreover, ΔCo-ac becomes null in the absence of previous correlated activity as reported experimentally (compare “TTX” in Fig. 6A and “naive” in Fig. 6B).

We use co-activation probability to investigate which parameters influence the distribution of synaptic clusters. We identified four parameters with which ΔCoac varies: (A) the input correlation (B) the number of compartments (C) the number of learning parameters (dimensionality of learning) and (D) non-linear summation within a compartment (four panels in Fig.7). While varying each parameter in turn, we kept all other parameters constant.

**Figure 6:**
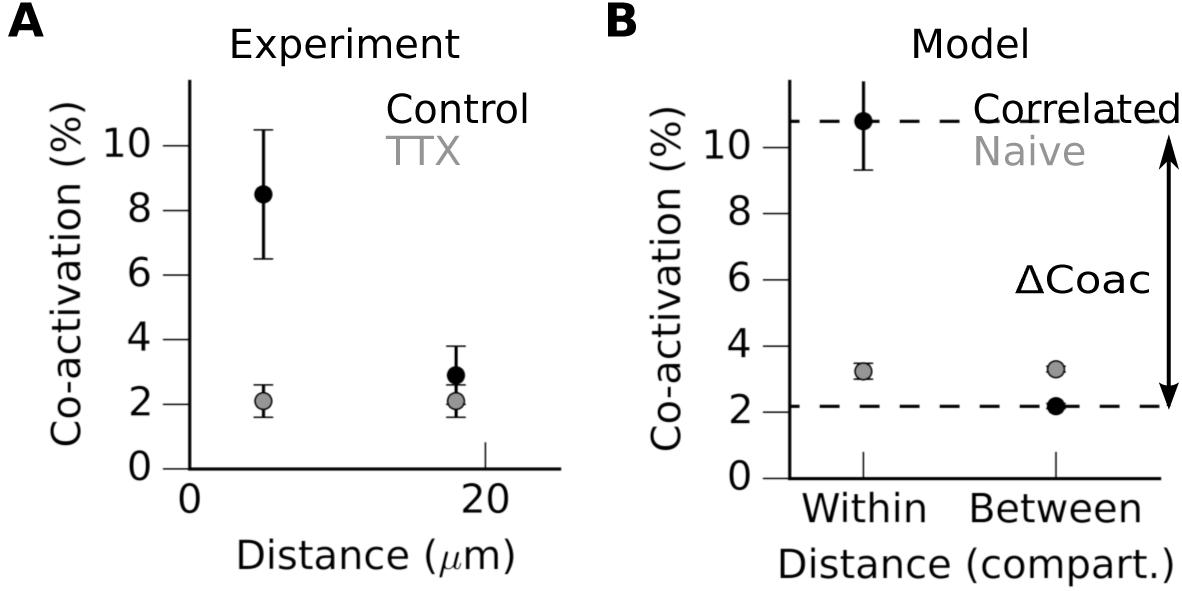
Co-activation probability quantifies synaptic cluster distribution. **A**.Measurements of spines co-activation probability in hippocampal slices, control situation and cultured within a TTX environment, washed out during measurement (replotted from Kleindienst et al 2011). **B**. Co-activation probability obtained in silico (see methods). Difference of co-activation “between” and “within” subunits, depending on the input correlation.

**Figure 7:**
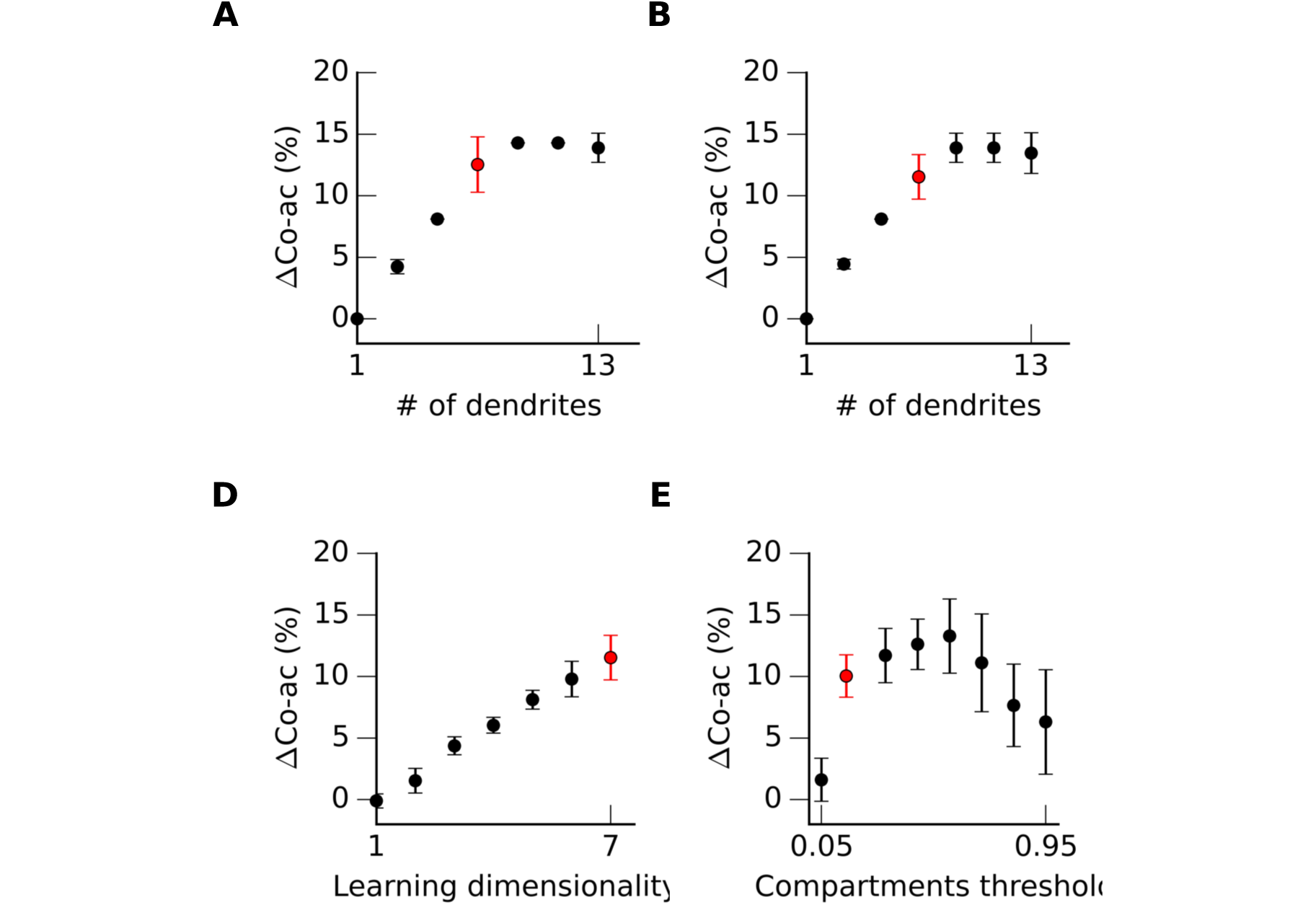
ΔCoac probability determines conditions favorable to the formation of synaptic clusters. ΔCoac is the difference between within and between co-activation probabilities. The mean ΔCoac probabilities (dots) are averaged over 100 model instances. We fix the value of all other parameters (red dot) while varying each in turn. We quantify the effect of **A** Input correlation within each ensemble; **B** The number of subunits in the neuron model; **C** Learning dimensionality which is the number of distinct values used in learning, e.g. a low learning dimensionality implies a high crosstalk between subunits; **D** the threshold (*θ*) within a non-linear subunit.

Correlation within an ensemble of input spikes must be sufficiently high to generate synaptic clustering. We found that in our model only input correlation greater or equal to 60% achieved a ΔCoac larger than 5% (Fig.7A). The more correlated the input, the higher the ΔCoac observed. We infer that correlations are necessary for the formation of synaptic clusters. Moreover, the formation of multiple synaptic cluster requires multiple compartments. Even in an integrate and fire model, synapses cluster in response to correlated inputs, and a single cluster forms. However, ΔCoac is always zero in that case, as all synapses target only one compartment. For example, in the case of seven input ensembles, three or more compartments are required for a ΔCoac larger than 5% (Fig.7B).

Even in a model with multiple compartments, it is possible to use a single *v* (e.g. membrane voltage or calcium concentration) that results from synaptic integration. In this case, *v* applies to all compartments and the learning dimensionality is one (i.e., maximum intra-compartmental cross-talk). Alternately, one can use multiple *v*s each assigned to distinct sets of compartments. For seven input ensembles, we found that three or more distinct vs are required for a ΔCoac larger than 5% (Fig.7C).

Finally, we studied the influence of the dendritic threshold on cluster distribution. Unlike the three other parameters, ΔCoac reaches its highest value in a large range (here between 0.1 and 0.4). For a high threshold (0.7 and above) integration is quasi-linear because synaptic activity does not reach the compartment’s threshold. Synaptic clusters can form even in this case (Fig.7D). Local non-linear integration is not necessary for synaptic cluster formation, but it guarantees their even distribution across compartments and novelty detection capacity.

## Discussion

Here we have demonstrated an unsupervised learning algorithm that generates synaptic clusters distributed evenly across compartments. This distribution is made possible by maintaining constant synaptic weights. Indeed, in our learning algorithm, each pre-synaptic neuron makes the same number of identical synaptic contacts on the postsynaptic neuron. Moreover, clusters organized in this way enable the neuron to differentiate novel from familiar stimuli, depending on dendritic and somatic spike thresholds.

We set a high somatic threshold requiring multiple dendritic spikes to trigger a somatic spike. In this case, the neuron detects novel stimuli and stays silent for familiar stimuli. Alternatively, a single dendritic spike might suffice to generate an action potential. Wu and Mel studied this scenario and described how dendrites could boost memory capacity at the network level (Wu and Mel, 2009). In this dual case, the neuron will fire for familiar stimuli and stay silent for novel stimuli and requires supra-linear summation in dendrites (Caze et al., 2012). Both cluster and scatter sensitivities could prevail depending on the cell type and the brain area.

A result of the learning scheme here is that neuronal ability to detect novelty will depend upon the intrinsic properties of each cell. A neuron with high dendritic threshold and low somatic threshold will detect familiar inputs, whereas a neuron with low dendritic threshold and high somatic threshold will detect novel inputs. Flexible computation is thus enabled by tuning cellular properties.

The learning algorithm presented here is unsupervised, but differs significantly from previous examples of this type of algorithm (Legenstein and Maass, 2011). Our work differs in two principle respects. First, we introduced horizontal normalization, in which an input makes a constant number of synapses on a post-synaptic neuron. This was initially introduced for biological realism as each neuron makes a limited number of synapses (Branco and Staras, 2009). This normalization turns out to avoid synaptic cluster repetition in multiple compartments of the post-synaptic neuron; with cluster repetition, detection of novel stimuli would be compromised. Second, rather than splitting inputs into training and test sets, we use a continuous learning paradigm. This renders our neuron model more realistic and enables it to continuously adapt to a changing sensory environment.

This computational work yields two immediate predictions: (1) A given neuron can display both clustered and scattered synaptic activity. This could be tested by *in vivo* two-photon imaging of the same neuron across periods of both spontaneous and sensory-evoked activity. (2) The number of synapses from a pre- to a post-synaptic neuron remains constant on short time scale. Time lapse imaging of synapses from a single afferent could test this prediction.

In conclusion, we have proposed a learning mechanism to organize the distribution of synapses in space. This mechanism could possibly underpin the emergence of stimulus selectivity underlying a wide range of neural computations.

## Acknowledgments

The author would like to thank Mark Humphries and Lyle Graham for their comments on earlier versions of the manuscript, and Jacopo Bono for proof reading of this manuscript.

## Supporting Information

### S1 Video

**Formation and distribution of synaptic clusters**. Activity is color-coded (white:inactive / black:active). The diameter of a circle relates to the number of spines on an input site. Triangle describes somatic activity.

### S2 Video

**Redistribution of synaptic clusters.**

### S3 Video

**Formation of synaptic clusters without horizontal normalization.**

### S4 Video

**Redistribution of synaptic clusters without horizontal normalization.**

### Software and code availability

We used Python v2.7, Numpy v1.3 and Matplotlib v1.4.0 to code, process and display the result of all our simulations. This code is available on a git repository (link to be supplied after acceptance of this manuscript, we uploaded this code for review with a README file).

